# MAIT cell responses to *S. aureus* and sensitivity to HlgAB are modulated by activation and tissue-dependent virulence effects

**DOI:** 10.64898/2026.03.17.712383

**Authors:** Elisa J. M. Raineri, Caroline Boulouis, Elli Mouchtaridi, Vera Nilsén, Curtis Cai, Tobias Kammann, Juliette Tabusse, Takuya Sekine, Nicole Wild, Christian Constantz, Eoghann White, Thomas R. Müller, Anne Marchalot, Sabrina Ferreira, Jyotsana Kaushal, Akhirunnesa Mily, Miriam Franklin, Elena Bonaiti, Mary-Lyn Eichhorn, John Bassett, Chris Stamper, Jeffrey Y. W. Mak, David P. Fairlie, Chris Tibbitt, Anna Norrby-Teglund, Nicole Marquardt, Jenny Mjösberg, Carl Jorns, Jenny Driving, Edwin Leeansyah, Marcus Buggert, Johan K. Sandberg

## Abstract

Mucosa-associated invariant T (MAIT) cells are unconventional T cells with innate-like rapid antimicrobial effector functions and serve as resident sentinels at mucosal and non-mucosal barriers. However, their role in immune defense against *Staphylococcus aureus* and the impact of bacterial immune evasion mechanisms are incompletely understood. Here, we have investigated MAIT cell responses to *S. aureus* and the impact of its broadly expressed leukocidin toxin HlgAB on MAIT cell responses in different human tissue sites. MAIT cells respond to *S. aureus* with a complex polyfunctional profile spanning pro-inflammatory IL-17, TNF, and IFNγ, anti-inflammatory IL-10, plus granzymes A, B, and K, perforin, and granulysin. The quality of responses was influenced by microbial dose and time of exposure and was dependent on both MR1-presented antigen and cytokine co-activation. CD56⁺ MAIT cells displayed stronger effector responses and higher HlgAB sensitivity compared to CD56⁻ cells. MAIT cells were partially resistant to HlgAB-toxicity compared to monocytes; blood-derived MAIT cells remained susceptible, whereas tonsillar MAIT cells showed minimal sensitivity. Notably, activation reduced the MAIT cell susceptibility to HlgAB, and such activation also afforded indirect protection to monocytes in co-cultures. The reduced susceptibility of tonsillar MAIT cells correlated with lower CCR2 and CXCR1 expression, a pattern shared with barrier tissues such as the lung and intestines. In conclusion, these findings indicate that MAIT cells exhibit tissue- and context-dependent responses to *S. aureus* and sensitivity to HlgAB-mediated immune evasion.

**Importance:** MAIT cells are an evolutionarily conserved unconventional T cell subset that responds to riboflavin pathway-derived antigens from a range of microbes. Here, we found that the human MAIT cell response to the pathogen *S. aureus* is robust with a polyfunctional complexity influenced by bacterial concentration and response kinetics. The ubiquitously expressed *S. aureus* immune-evasive toxin HlgAB attacks MAIT cells via CCR2. However, the sensitivity of MAIT cells to HlgAB varies depending on tissue localization, where in particular tissue-resident MAIT cells in tonsils are resistant. Antigen-specific activation of MAIT cells reduces HlgAB sensitivity, with protection also afforded to monocytes in the vicinity. These findings uncover the complex and dynamic interaction between an evolutionarily conserved arm of immunity, and immune evasion mechanisms of the important pathogen *S. aureus*.

## Introduction

Mucosa-associated invariant T (MAIT) cells are unconventional αβ T cells with rapid innate-like response characteristics and contribute to immune control of a range of infections (1–3). MAIT cells recognize antigens presented by the conserved and largely non-polymorphic major histocompatibility complex (MHC) class I-related (MR) 1 molecules (4, 5). Derivatives of unstable riboflavin biosynthesis pathway intermediates represent the main class of MR1 presented antigen and are expressed by many microbes (4, 5), including *Staphylococcus aureus*. MAIT cells are tissue-resident cells and are found abundantly throughout the human body (6), and enriched in barrier tissues and the liver (6–9). MAIT cells activated by MR1-presented antigens provide effector functions by rapid cytokine production (7, 10) and mediate strong cytolytic function against infected cells (11–13) and free-living bacteria (14). In addition, the MAIT cell response can be activated by cytokines including IL-12, IL-18 and type I IFNs, which enhances TCR-mediated activation (15–17) and triggers MR1-independent responses (18–20). Thus, MAIT cells are poised to respond to infection from a variety of pathogens and influence disease outcomes. Despite their evolutionary conservation and limited TCR diversity, MAIT cells display phenotypic and functional heterogeneity, with microbe-specific and subset-dependent specialization of effector responses (6, 20, 21). Emerging evidence from both humans and mice indicate that this phenotypic plasticity is reflected in the functional profiles of MAIT cells which are anatomically distinct and shaped by the local tissue microenvironment (22, 23).

*Staphylococcus aureus* is a leading cause of infections worldwide due to its ability to evade immune defenses and develop resistance to antibiotics (24–26). These gram-positive bacteria exemplify the spectrum of host-pathogen interactions, which include transitioning between commensal, colonizing, or invasive characteristics depending on the dynamic interplay with the host (27, 28). Evasion of the host immune response is crucial for the ability of *S. aureus* to establish infection, and it employs a wide range of virulence factors, with cytolytic toxin production being a primary mechanism (27, 29). The leukocidins belong to a family of bicomponent pore-forming toxins which lyse leukocytes and red blood cells (30, 31). This family includes Panton-Valentine Leukocidin LukSF (PVL), γ-haemolysin (HlgAB and HlgCB), LukED, and LukAB (also referred to as LukGH). The γ-haemolysin genes are part of the core genome found in more than 99.5% of human *S. aureus* isolates, whereas the other leukocidins, such as PVL and LukED, are considerably less prevalent. These genes can be gained or lost through horizontal gene transfer and are not essential for basic bacterial survival, as evidenced by their lack of universal conservation (32–36). The leukocidin receptor expression patterns enable the toxins to selectively and efficiently target both phagocytic cells and other leukocyte populations by targeting G-protein coupled receptors. For instance, LukED targets CCR5, CXCR1 and CXCR2. Interestingly, MAIT cells express high levels of CCR5 and are hypersensitive to LukED-mediated lysis (37). However, HlgAB targets CCR2, CXCR1 and CXCR2 and contributes to *S. aureus* bacteremia via CCR2-mediated mechanisms (34).

MAIT cell involvement in *S. aureus* immunopathogenesis remains incompletely characterized with only a few studies focused on peripheral blood MAIT cells rather than tissue-resident populations (37–40). In this study, we investigate MAIT cell interactions with *S. aureus* using high-parameter techniques and a range of human tissue samples, including peripheral blood, palatine tonsils, and matched mucosal and lymphoid tissues from organ donors. We specifically focus on the immune-evasive virulence factor HlgAB, which has near universal prevalence and consistent toxin expression across various *S. aureus* strains. We demonstrate that MAIT cells display tissue- and context-dependent phenotypic and functional heterogeneity, which shapes their response to *S. aureus* and its immune-evasive toxins. These findings further substantiate the complexity underlying *S. aureus* leukocidin-target cell interactions and highlight that MAIT cells represent an important underappreciated tissue-resident population with a unique susceptibility and response profile to *S. aureus*.

## Results

### MAIT cells exhibit a time- and dose-dependent polyfunctional response to *S. aureus*

MAIT cells are known to respond to *S. aureus in vitro* with rapid upregulation of CD69 and expression of cytokines (37, 39). To further investigate the polyfunctional profile and kinetics of MAIT cell responses, we pulsed THP-1 cells with mildly fixed *S. aureus* for 3 h and used these to stimulate MAIT cells isolated from PBMCs (Fig. 1A, Supplementary Fig. 1A). MAIT cells expressed a broad cytokine profile, including proinflammatory TNF and IFNγ and anti-inflammatory IL-10, along with cytotoxic mediators, including perforin, CD107a, granzyme (Gzm) A, B, and K, perforin, and granulysin (Fig. 1B-D). Expression of IL-17 and IL-22 was relatively low, reflecting the functional characteristics of blood MAIT cells.

**Figure 1.**
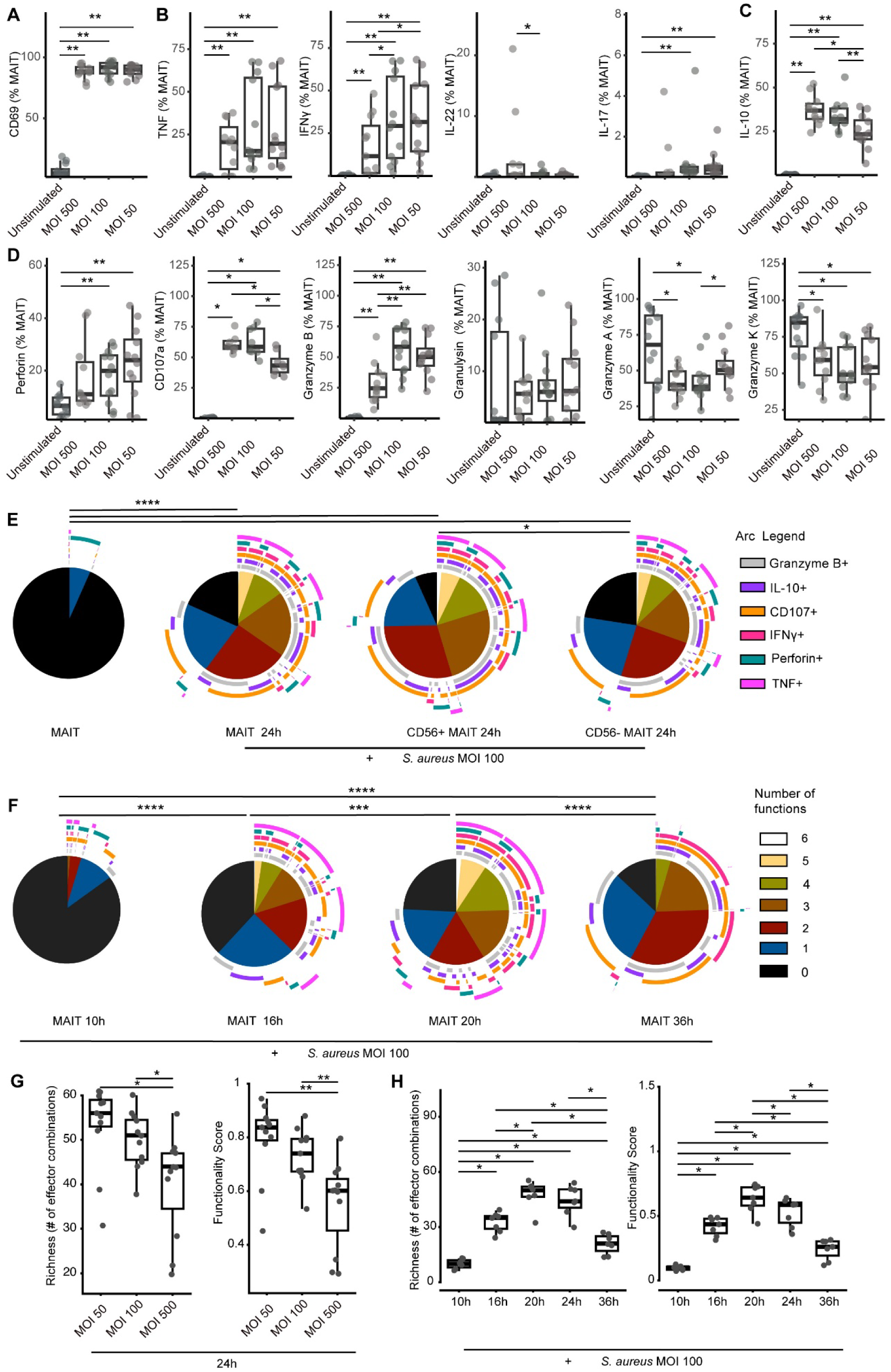
The complex polyfunctional MAIT cell response to *S. aureus*. (A) Peripheral blood MAIT cells response to *S. aureus*-pulsed THP-1 cells assessed after 24 h with the proxy marker for cell activation CD69 for different MOI. MAIT cells were analysed for a broad cytokine profile, spanning from (B) pro-inflammatory TNF, and IFNγ, IL-22, and IL-17, to (C) anti-inflammatory IL-10, along with cytotoxic mediators including (D) perforin, CD107a, granzymes A, B, and K, perforin, and granulysin. (E) The polyfunctionality profile of all MAIT cells stimulated with *S. aureus*-pulsed THP-1 cells for 24 h or separated in CD56+ and CD56-MAIT cell subsets, in terms of numbers of functions (pie slice) and the type of cytokine (arc) (n=9). (F) Pie charts displaying temporal changes in MAIT cell polyfunctionality following *S. aureus* stimulation compared with the unstimulated condition at 10 h, 16 h, 20 h and 36 h (n=9). (G, H) Analysis of polyfunctionality using the COMPASS algorithm. Richness of the immune response quality and functionality score indicating the proportion of responding effector subsets among all possible Boolean combinations across different MOI at 24 h and across time at MOI 100. Statistical comparison between conditions with the Wilcoxon matched-pairs signed-rank test for (A, B, C, D, G, H), p-values were adjusted using the Benjamini-Hochberg method. Statistical comparison between the MAIT cell populations with the permutation test (E and F). *p<0.05, **p<0.01, ***p<0.001 and ****p<0.0001.”

The MAIT cell response magnitude for different functions varied across graded multiplicities of infection (MOI). IFNγ and cytolytic effector molecules perforin and GzmB showed peak expression at MOI 100 or 50, whereas CD107a and IL-10 were higher at MOI 500 (Fig. 1B-D). Thus, the MAIT cell response to *S. aureus* shifts depending on the bacterial dose, from predominantly IFNγ and TNF at lower bacterial concentrations towards a profile dominated by cytolytic degranulation and IL-10 expression at higher bacterial concentrations.

The MAIT cell response to *S. aureus* displayed a complex polyfunctionality pattern with diverse combinations of one to six functions co-expressed by 24 h of stimulation (Fig. 1E). Stratification of MAIT cells by CD56 expression revealed that CD56^+^ MAIT cells displayed significantly greater polyfunctionality in the response against *S. aureus* than cells negative for CD56 (Fig. 1E). This finding is consistent with previous studies demonstrating that CD56^+^ MAIT cells have an enhanced capacity to respond to the innate cytokines IL-12 and IL-18 and overall enhanced effector capacity (6, 20).

We next evaluated the polyfunctional kinetics of MAIT cells responding to *S. aureus* at 10, 16, 20 and 36 h. Limited mono- or dual-functional responses were detected at 10 h, complexity increased progressively at 16 h, peaking at 20 h and 24 h, before decreasing at 36 h (Fig. 1F). TNF expression was particularly low at the 36 h time point. Altogether, these findings indicate that the quality of the MAIT cell response to *S. aureus* evolves over time and is influenced by microbial dose. To gain deeper insight into the quality of the MAIT cell-mediated immune response, we applied the Combinatorial Polyfunctionality Analysis of Antigen-Specific T Cell Subsets (COMPASS) framework. Briefly, COMPASS employs a Bayesian hierarchical model to identify functional subsets, benchmarked against donor-matched unstimulated control cells. At 24 h, COMPASS analysis indicated that MAIT cell polyfunctionality was inversely related to bacterial dose, with both functionality score and response richness being highest at MOI 50 and progressively declining at MOI 100 and MOI 500 (Fig. 1G). This pattern suggests that high bacterial loads impair rather than enhance MAIT cell functional effector diversity. In the time course analysis at MOI 100, MAIT cell polyfunctionality varied across time points, with peak levels at 20 h as indicated by the highest functionality score and response richness (Fig. 1H and Supplementary Fig. 1B).

### MAIT cell activation in response to *S. aureus* depends on both MR1 and innate cytokines

To investigate the primary mode of MAIT cell activation by *S. aureus*, blocking antibodies against MR1 and innate cytokines were added to co-cultures of isolated Vα7.2^+^ cells with *S. aureus*-pulsed THP-1 (Fig. 2). Combined blockade of IL-12 and IL-18 largely abolished the production of the proinflammatory cytokines TNF and IFNγ. MR1 blockade also reduced expression of these cytokines, albeit to a lesser extent. A similar trend was observed for the induction of GzmB expression. MR1 blockade had a more pronounced inhibitory effect on the expression of IL-10, granulysin and CD107a degranulation compared to innate cytokines blockade. Bacterial stimulation led to a significant decrease in intracellular GzmA and GzmK levels, consistent with degranulation and release by the responding MAIT cells. However, MR1 blockade mainly restored intracellular GzmK levels, whereas GzmA restoration was observed with both MR1 and cytokine blockade.

**Figure 2.**
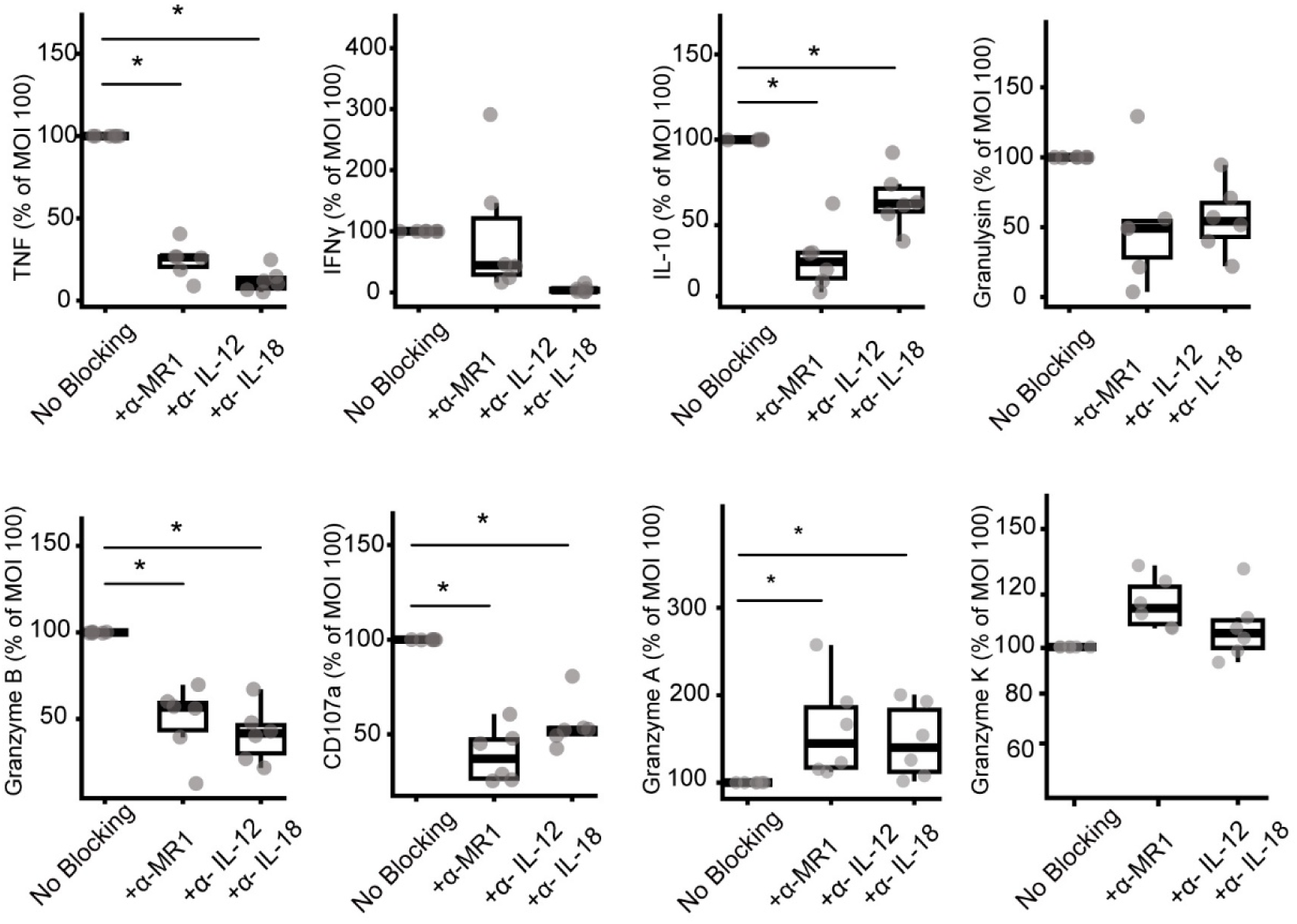
MAIT cell activation in response to *S. aureus* depends on both MR1 and innate cytokines. MAIT cell response to *S. aureus* at MOI 100 dependency on both MR1-presented antigen and cytokine co-activation as assessed by blocking with mAbs. MAIT cells were analysed for a broad cytokine profile, spanning from pro-inflammatory TNF, and IFNγ to anti-inflammatory IL-10, along with cytotoxic mediators including granulysin, granzyme B, CD107a, granzyme A and granzyme K. Statistical comparison between conditions with the Wilcoxon matched-pairs signed-rank test, p-values were adjusted using the Benjamini-Hochberg method. *p<0.05, **p<0.01, ***p<0.001 and ****p<0.0001.

### Leukocyte subset sensitivity to HlgAB cytotoxicity

To examine the impact of HlgAB toxin on cells in peripheral blood, PBMCs were incubated with recombinant HlgA and HlgB proteins and analysed for population changes by flow cytometry, using Uniform Manifold Approximation and Projection (UMAP) analysis. The defining markers (CD3, CD4, CD8, CD161, Vα7.2, CD56, CCR2, CXCR1, CXCR2 and CD14) were projected onto the UMAP to identify distinct cell populations such as T cells, MAIT cells and monocytes (Fig. 3A). UMAP projection of HlgAB-treated versus untreated conditions revealed differences in cellular composition and the loss of specific populations. The UMAP topography corresponding to CD14^+^ cells was nearly absent following HlgAB exposure, suggesting that the toxin depletes monocytes (Fig. 3B). The region defined by Vα7.2 and CD161 was also impacted by toxin treatment, albeit to a lesser extent, whereas CD3^+^ non-MAIT T cells were less affected (Fig. 3B). These differences were more evident when different concentrations of toxin were compared: At a lower HlgAB concentration of 0.31 µg/ml, we observed approximately 25% survival of CD14⁺ cells, around 75% survival of MAIT cells, and nearly complete survival (close to 100%) of non-MAIT T cells, compared to the untreated sample (Fig. 3C). Higher doses of 2.5 µg/ml and 10 µg/ml led to almost complete loss of both MAIT cells and CD14^+^ cells. In contrast, non-MAIT T cells showed 25% survival at 2.5 µg/ml, whereas exposure to the highest HlgAB dose resulted in essentially complete T cell death. Division of MAIT cells based on CD56 expression, showed a differential sensitivity to HlgAB exposure at lower toxin doses, with approximately 75% and 50% survival of CD56^-^ and CD56^+^ MAIT cells, respectively.

**Figure 3.**
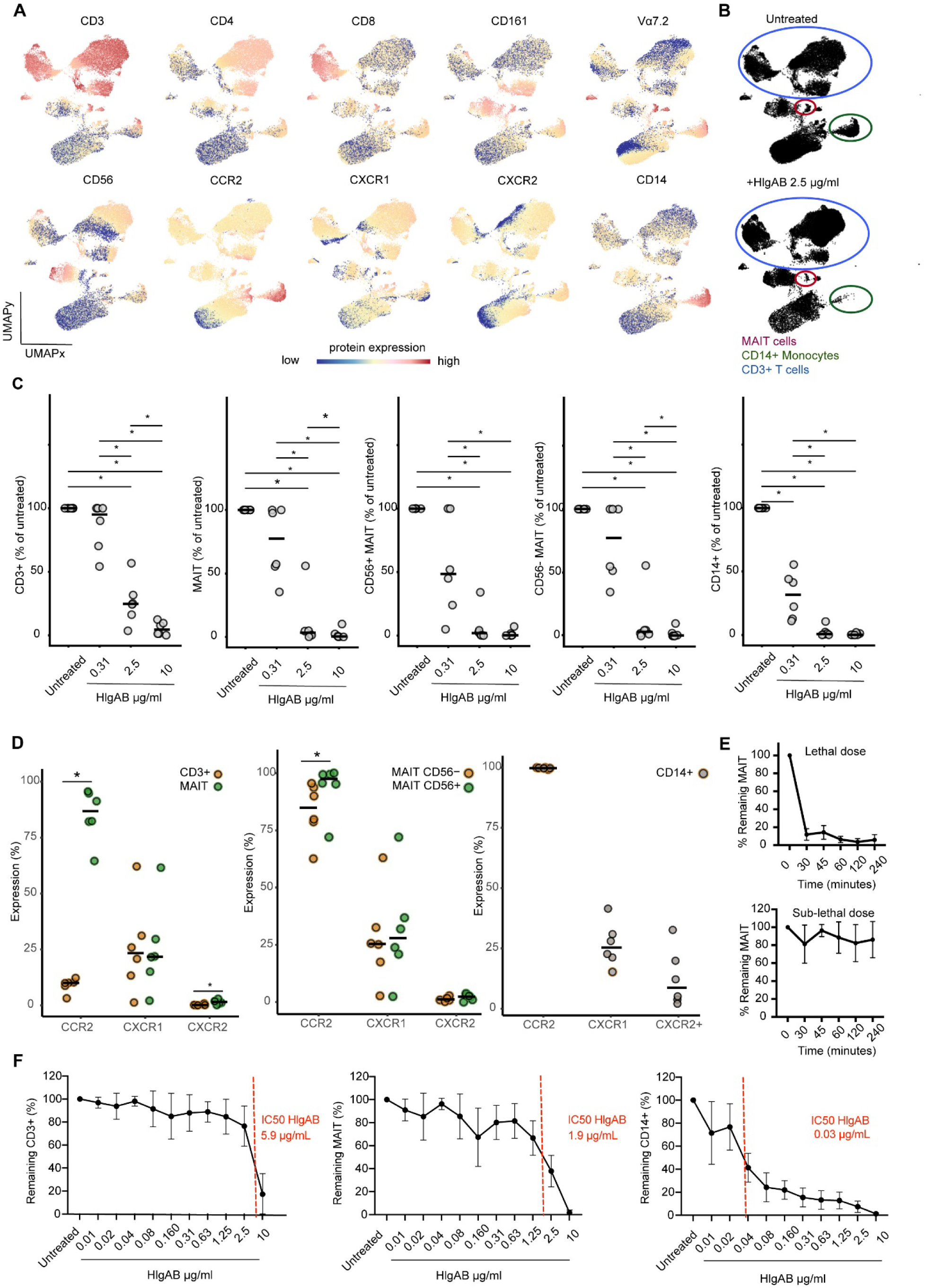
Differential susceptibility to HlgAB among leukocyte subsets, with monocytes being the most susceptible and MAIT cells showing intermediate resistance. (A) UMAP plots of total live PBMCs showing expression of the indicated markers. (B) UMAP plots of total PBMCs exposed to HlgAB for 1 h or not (n=6). (C) Relative percentages of total CD3^+^ cells, MAIT cells, CD56^+^ and CD56^-^ MAIT cells, and CD14^+^ cells after exposure to a range of HlgAB concentrations for 1 h. (D) CCR2, CXCR1 and CXCR2 expression across CD3^+^ cells, MAIT cells, CD56^+^ and CD56^-^ MAIT cells, and CD14^+^ cells. (E) Relative percentages of MAIT cells after exposure to a high dose (10 µg/ml) and low dose (0.01 µg/ml) of HlgAB during a time course from 30 min to 4 h (n=5). (F) Relative percentages of CD3^+^ cells, MAIT cells and CD14^+^ cells over a range of HlgAB concentrations after 1 h of exposure. The IC50 is indicated in red (n=5). Statistical comparison between conditions with the Wilcoxon matched-pairs signed-rank test (C-D) and p-values were adjusted using the Benjamini-Hochberg method (C). Statistical comparison between conditions with the Wilcoxon matched-pairs signed-rank test for (C and D). *p<0.05, **p<0.01 and ***p<0.001.

Since it was previously shown that the human chemokine receptors CCR2, CXCR1 and CXCR2 act as receptors for HlgAB, we analysed the expression of these receptors on cell subsets in blood. The region defined by CD14^+^ on the UMAP topography had the highest expression of CCR2 (Fig. 3A), and this was confirmed by the 100% expression level on gated CD14^+^ cells (Fig. 3D). The vast majority of MAIT cells expressed CCR2, in stark contrast to non-MAIT T cells, which displayed very low CCR2 expression (Fig. 3D). CD56^+^ MAIT cells expressed more CCR2 than CD56^-^ MAIT cells, which might explain the higher sensitivity to HlgAB detected at lower doses (Fig. 3D). Furthermore, approximately 25% of non-MAIT T cells, MAIT cells and CD14^+^ cells expressed CXCR1. In contrast, CXCR2 expression was absent from non-MAIT T cells and MAIT cells, while CD14^+^ cells showed low expression of CXCR2.

We next examined the kinetics and dose-dependency of HlgAB toxicity (Fig. 3E and 3F). MAIT cell depletion occurred rapidly, within 30 min of toxin exposure (Fig. 3E). Notably, MAIT cells remained viable at sub-lethal doses over time, indicating that prolonged exposure did not further enhance cell depletion (Fig. 3E). Furthermore, the IC_50_ of HlgAB was lower for CD14^+^ cells (0.03 µg/ml) than for non-MAIT T cells (5.9 µg/ml) and MAIT cells (1.9 µg/ml) (Fig. 3F, red reference line). Thus, 60-fold higher doses of toxin are needed to deplete MAIT cells as compared to monocytes.

We next evaluated the small molecule CCR2 antagonists, BMS CCR2 22 and cenicriviroc (CVC), for their ability to protect MAIT cells from HlgAB cytotoxicity (Supplementary Fig. 2). BMS CCR2 22 features a *cis*-1,2-diaminocyclohexane scaffold and represents a new class of CC chemokine receptor 2 antagonists (41). Cenicriviroc is a dual CCR2/CCR5 antagonist, previously studied in clinical trials for the treatment of HIV or non-alcoholic steatohepatitis (NASH), that is also of interest for other inflammatory conditions (42, 43). The addition of BMS CCR2 22 did not rescue the MAIT cell population from HlgAB cytotoxicity (Supplementary Fig. 2A), and neither did addition of CVC rescue MAIT cells or monocytes (Supplementary Fig. 2B-C). This could be due to different binding sites for CCR2 antagonists and HlgAB (44, 45). Alternatively, residual toxin binding to CXCR1, despite its low expression on MAIT cells, might contribute to the incomplete blockade. However, further structural and functional experiments are needed to clarify the reason for these findings.

### TCR-mediated activation diminishes HlgAB sensitivity both in MAIT cells and in surrounding antigen-presenting monocytes

MAIT cells are activated by TCR recognition of MR1-presented microbial vitamin B2 metabolite derivatives such as 5-OP-RU. TCR-independent activation can occur by innate cytokines such as IL-12 and IL-18 primarily released by APCs during infection or inflammation (2). To examine whether MAIT cell activation would change the outcome of HlgAB exposure, we stimulated MAIT cells with 5-OP-RU or with a combination of IL-12 and IL-18 (Fig. 4A, Supplementary Fig. 3). Notably, 5-OP-RU stimulated samples showed preservation of monocytes compared to unstimulated samples after HlgAB exposure (Fig. 4A and B). Similarly, MAIT cells exhibited a trend toward higher survival relative to unstimulated samples, a pattern consistent in both CD56^+^ and CD56^-^ subsets. This effect coincided with decreased CCR2 and CXCR1 expression on MAIT cells in stimulated samples (Fig. 4C), whereas no significant changes were detected in CCR2 and CXCR1 expression on CD14^+^ cells (Fig. 4C). Taken together, these observations suggest that MAIT cell TCR activation partly prevents HlgAB-induced cell loss, both in MAIT cells themselves and in monocytes in the microenvironment. This might be a mechanism of protection in an environment with high bacterial density and the concomitant availability of vitamin B2 metabolite-derived antigen.

**Figure 4.**
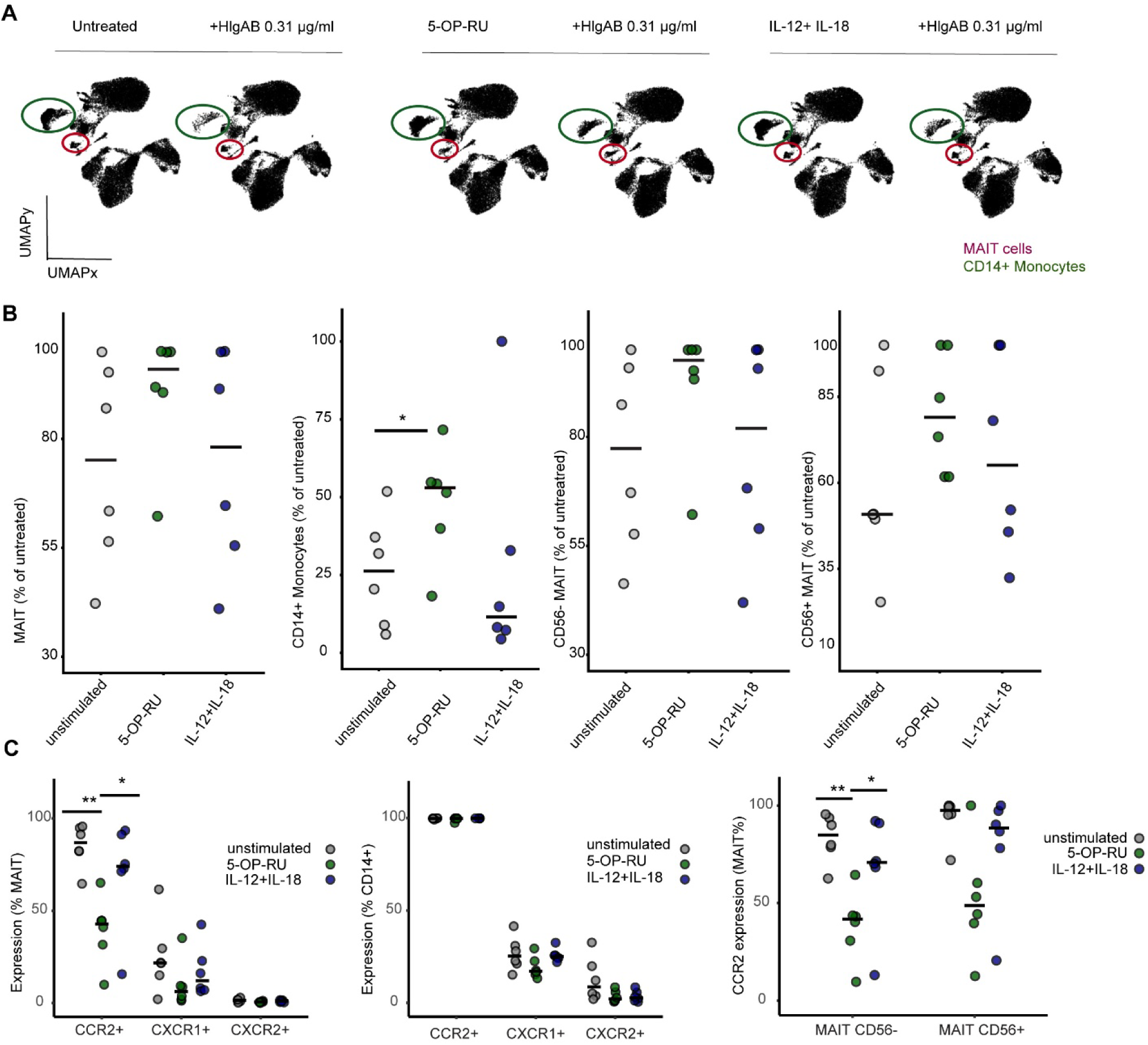
MAIT cell TCR activation drives decreased sensitivity to HlgAB, both in MAIT cells themselves and in CD14⁺ monocytes. Total PBMCs under unstimulated conditions or following TCR and innate cytokine stimulation, in the presence or absence of HlgAB (0.31 µg/ml) for 1 h. The corresponding UMAP plots of total live PBMCs showing expression of the indicated markers is shown in Supplementary Figure 3. (A) UMAP visualization of PBMCs showing alterations in MAIT cell and CD14⁺ monocyte compartments following HlgAB-induced cytotoxicity (n=6). (B) MAIT cells, CD56⁺ and CD56⁻ MAIT cells, and CD14⁺ monocytes after 1 h exposure to HlgAB, plotted as % of the respective untreated control. (C) Expression of CCR2, CXCR1, and CXCR2 across MAIT cells, CD56⁺ and CD56⁻ MAIT subsets, and CD14⁺ monocytes under stimulated and unstimulated conditions. Statistical comparison between conditions with the Wilcoxon matched-pairs signed-rank test (B and C). *p<0.05, **p<0.01, and ***p<0.001.

### Tissue localization influences expression of leukocidin receptors on MAIT cells

To evaluate if the tissue localization can affect the expression of leukocidin receptors on resident MAIT cells, we obtained matched samples from blood, spleen, liver, ileum, caecum, colon and lung, as well as the associated lymph nodes (LN) from lung and mesenterium, from nine human organ donors, and analyzed the isolated cells by flow cytometry. Tissue MAIT cells were identified as cells expressing CD3, CD45, CD161, Vα7.2, and staining positive with the MR1 tetramer loaded with 5-OP-RU (Fig. 5A). MAIT cell frequencies showed significant donor variability, and a tissue-dependent pattern with significantly higher frequencies in liver compared to peripheral blood and reduced in the lung LNs (Fig. 5B). This corroborates our previous studies indicating tissue- and donor-dependent patterns in the human MAIT cell compartment (6). To gain deeper insight into the MAIT cell compartment in tissues, we examined the expression of CD69 and CD56 (Fig. 5C and D). CD69 expression was high in all tissues compared to blood supporting the interpretation that these cells exhibit features of tissue residency. In contrast, CD56 expression varied between tissues and was particularly elevated in the spleen and in the liver.

**Figure 5.**
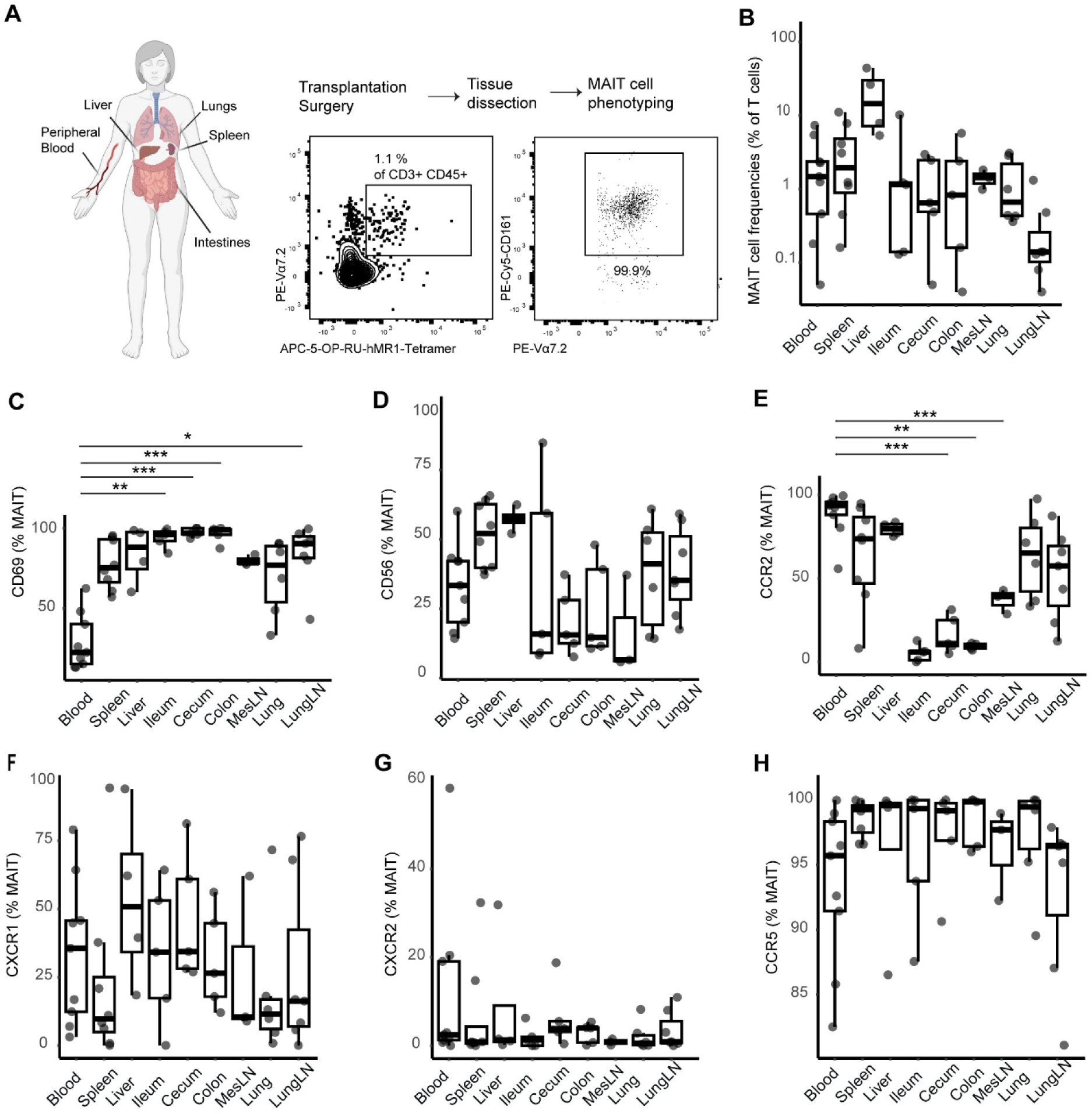
Heterogenous expression of leukocidin receptors in human tissues. (A) Tissue samples were obtained from human organ donors and MAIT cells were subsequently identified as live CD45^+^, CD3^+^ 5-OP-RU-MR1-tetramer^+^ Vα7.2^+^ CD161^+^ lymphocytes. (B) MAIT cell frequency was calculated as the percentage of T cells for each donor and tissue and is shown on a logarithmic scale. Surface expression of (C) CD69 and (D) CD56 by MAIT cells. (E) Surface expression of leukocidin receptors CCR2, CXCR1, CXCR2 and CCR5 by total MAIT cells. The number of samples analyzed (donors=9): blood (n=9), spleen, (n=8), liver (n=4), ileum (n=5), cecum (n=5), colon (n=5), mLN (n=3), lung (n=6) and lungLN (n=7). In each tissue site, only samples with more than 20 gated MAIT cells were included in the phenotypic analysis. Differences in MAIT cell frequencies or marker expression between blood and each tissue were analyzed using Kruskal-Wallis test followed by Dunn’s post-hoc test for all pairwise comparisons. *p<0.05, **p<0.01, ***p<0.001 and ****p<0.0001.

Given that *S. aureus* establishes infection within tissues before disseminating into the bloodstream, we assessed the expression of the HlgAB receptors CCR2, CXCR1 and CXCR2 in tissue-resident MAIT cells. In this analysis, we also included the receptors for LukED considering this toxin targets both CCR5 and CXCR1/CXCR2. We observed significant differences in MAIT cell CCR2 expression in tissues compared to blood (Fig. 5E). CCR2 expression was nearly undetectable in the ileum, colon, and caecum, while mesenteric LNs exhibited low expression, though not as low as intestinal tissues. CCR2 levels were low in MAIT cells isolated from lungs and lung LNs when compared to blood, whereas it was higher in the liver. CXCR1 was expressed by a minority of MAIT cells across all tissues, with no significant differences between tissues (Fig. 5F). Additionally, while CXCR2 was mostly absent on MAIT cells across tissues (Fig. 5G). In contrast to CCR2, the LukED receptor CCR5 was highly expressed on MAIT cells in all tissues with the highest expression in the spleen (Fig. 5H). Altogether, these results indicate differential expression of CCR2 and CCR5 in MAIT cells resident across different tissues, suggesting that MAIT cell sensitivity and responses to *S. aureus* leukocidins are tissue site dependent.

### Tonsillar MAIT cells exhibit reduced susceptibility to HlgAB and diminished CCR2 expression

*S. aureus* is one of the most frequent pathogens in tonsillitis (46, 47), and a major cause of recurrent tonsillitis, showing intracellular persistence and the ability to form multilayered biofilms on tonsillar tissues, the latter likely contributing to its persistence in chronic infections (46, 48, 49). In tonsils, MAIT cells are found near germinal centers contributing to both the innate antimicrobial response and adaptive B cell help (50). Tonsillar MAIT cells were identified as cells expressing CD3, CD45, CD161, Vα7.2, and staining positive with the MR1 tetramer loaded with 5-OP-RU (Fig. 6A). We investigated tonsils (n=32) and matched blood (n=7) and first assessed overall T cell frequencies in blood versus tonsillar tissues, revealing a significant enrichment of CD4^+^ T cells and lower frequency of CD8^+^ T cells in the tonsil compared to blood (Fig. 6B). MAIT cell frequencies were approximately six-fold lower in tonsil than in blood, and CD56 expression was substantially reduced in tonsillar MAIT cells (Fig. 6C).

**Figure 6.**
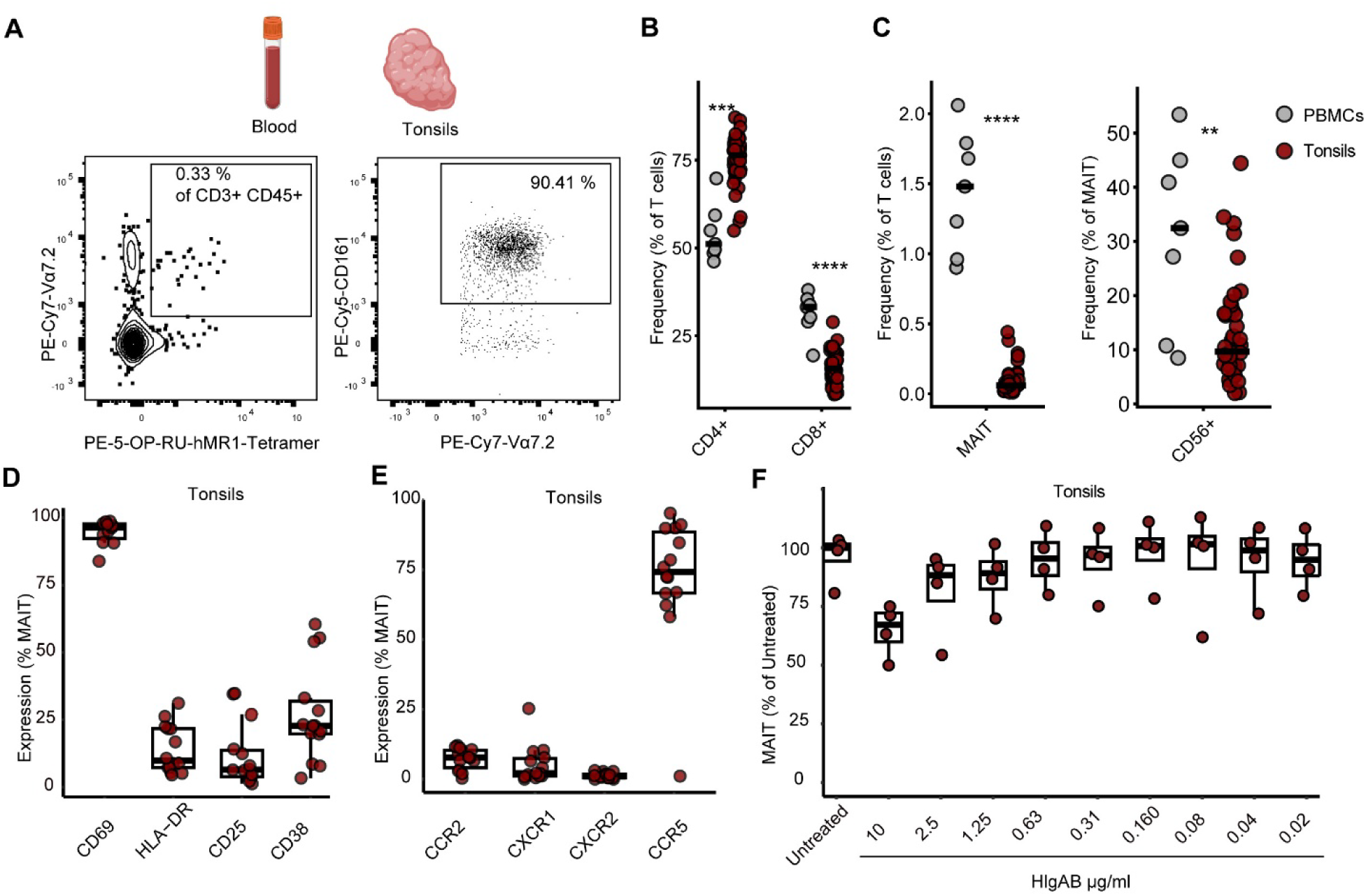
MAIT cells in the tonsils are relatively resistant to HlgAB, a feature associated with lower leukocidin receptor expression. Peripheral blood and tissue samples were obtained from individuals undergoing tonsillectomy for recurrent infections or obstructive sleep apnea. (A) MAIT cells were identified as live CD45^+^ CD3^+^ 5-OP-RU-MR1-tetramer^+^ Vα7.2^+^ CD161^+^ lymphocytes. (B) Frequencies of total T cells and their CD8⁺ and CD4⁺ subsets in peripheral blood versus tonsils, and (C) frequencies of MAIT cells, including CD56⁺ and CD56⁻ subsets, in peripheral blood versus tonsils (blood n=7 and tonsils n=25). (D) Expression of activation markers CD69, HLA-DR, CD25 and CD38 on tonsillar MAIT cells to assess their activation status (n=14). (E) Expression of CCR2, CXCR1, CXCR2 and CCR5 across tonsillar MAIT cells (n=14). (F) Relative percentages of tonsillar MAIT cells over a range of HlgAB concentrations after 1 h of exposure (n=4). Statistical comparisons between PBMCs and tonsillar cells were performed using the Wilcoxon rank-sum test (B, C). Comparisons between tonsillar MAIT cells under different treatment conditions were performed using the Wilcoxon matched-pairs signed-rank test (F). *p<0.05, **p<0.01, ***p<0.001 and ****p<0.0001.

To further characterize the MAIT cell phenotype in the tonsillar environment we evaluated the expression of CD69, HLA-DR, CD25 and CD38 on fresh tonsil samples (n=14) (Fig. 6D). Tonsillar MAIT cells were CD69+, supporting the interpretation that these cells exhibit a tissue-resident phenotype. The classical activation markers HLA-DR, CD25 and CD38 remained low to intermediate in the tonsils. Expression of leukocidin receptors was low, with the exception of CCR5, which was expressed on approximately 75% of tonsillar MAIT cells but remained lower than expression levels observed in healthy blood and organ donor tissues (Fig. 6E). We next investigated whether the low expression of CCR2 and CXCR1 in tonsillar MAIT cells affected their toxin sensitivity. Tonsillar cells were incubated with the same concentration range previously tested in blood (Fig. 6F). Unlike blood-derived MAIT cells (Fig. 3), tonsillar MAIT cells showed no significant loss of viability, even at higher concentrations, indicating a marked resistance to HlgAB-mediated toxicity. Altogether, these findings indicate that MAIT cells exhibit tissue- and context-dependent phenotypic and functional diversity, which shapes their sensitivity to *S. aureus* immune-evasive toxins.

## Discussion

The interactions between *S. aureus* and the immune system are complex, and despite more than a century of research, this bacterium is still a leading cause of invasive infections. Moreover, the increasing incidence of multidrug-resistant *S. aureus*, particularly methicillin-resistant *S. aureus*, poses a major health threat. Even though there is a growing interest in the role of MAIT cells due to their therapeutic potential, little is known about their contribution to *S. aureus* immunity and pathogenesis and their possible role in antimicrobial therapy (39, 51, 52). MAIT cells use diverse mechanisms to control bacterial infections, prevent tissue damage and synergize with antibiotics to kill bacteria (51). They also directly kill bacterially infected cells via rapid GzmB and perforin induction in an MR1-dependent manner (13, 14). In this study, we show that MAIT cells respond to *S. aureus* with a broad polyfunctional cytokine response including TNF, IFNγ, IL-10, as well as production of cytotoxic molecules GzmA, B, K, perforin, and granulysin. This functional diversity was particularly pronounced in CD56⁺ MAIT cells, which displayed significantly greater polyfunctionality than CD56⁻ MAIT cells. Characterization of MAIT cell response kinetics revealed that duration of the response was a critical determinant of this polyfunctionality. These findings highlight the rapid activation and diverse effector functions of MAIT cells in response to *S. aureus,* involving MR1-restricted pathways and signalling via innate cytokines. Notably, MAIT cell effector responses show a high degree of plasticity; recent findings indicate that they can adapt these effector responses based on cytokines in the microenvironment (22). For instance, IL-23 was reported to co-stimulate antigen-specific MAIT cell activation enabling control of pulmonary *Legionella* infection in mice (53). In an infection model of chronic *Mycobacterium tuberculosis* infection, 5-OP-RU stimulation supported MAIT cell expansion and an IL-17A-dependent reduction in bacterial loads (54). Interestingly, MAIT cells can be expanded *in vitro* with maintained characteristics and enhanced cytotoxic potential (55). Further studies may be designed to investigate whether MAIT cell responses could be utilized in combination with antibiotics as therapeutic mediators to enhance control of antimicrobial-resistant *S. aureus* infections.

The leukocidin bi-component pore-forming family includes PVL (targeting C5aR1/ C5aR2), LukED (targeting CCR5/CXCR1/CXCR2), HlgAB (CCR2/CXCR1/CXCR2), HlgCB (targeting C5aR1/C5aR2) and LukAB (targeting CD11b) (30). The receptor specificities of leukocidins create a variable landscape of cellular vulnerability. MAIT cells represent an underestimated target T cell population in light of their unique position as a tissue-resident population with mixed innate and adaptive immune characteristics and a distinct receptor profile. It was previously shown that MAIT cells are hypersensitive to CCR5-dependent LukED toxicity compared to conventional T cells (37). Here, we focused on MAIT cell sensitivity to HlgAB, a critical virulence factor due to its near universal prevalence and consistent expression by various *S. aureus* strains. We observed cell type-specific sensitivity to HIgAB within human PBMC populations with MAIT cells being more resistant to toxin-induced cell death than monocytes, yet more susceptible than conventional T cells. Furthermore, the relative resistance to HlgAB observed in CD56^-^ MAIT cells was reflected by a lower CCR2 expression as compared to their CD56^+^ counterparts. This finding may be important as CD56 expression is associated with elevated effector functions in MAIT cells, a phenotype enriched in the liver (6), which may render the hepatic microenvironment favourable for *S. aureus* regarding immuno-evasion.

CCR2 was universally expressed on CD14^+^ cells, relatively highly expressed on MAIT cells, and with low expression on non-MAIT T cells. CXCR1 was minimally expressed across all three cell types, while CXCR2 was absent from both T cell populations and with only minimal expression on CD14^+^ cells. Interestingly, within the PBMC pool TCR-mediated MAIT cell activation shifted HlgAB mediated toxicity towards increased cellular resistance and survival, with reduced toxicity both against MAIT cells themselves and against monocytes. This observation suggests a protective mechanism mediated by MAIT cells and adds context to the associated downregulation of CCR2 expression.

The sensitivity of MAIT cells to HlgAB toxicity displayed substantial variation depending on tissue site. Tonsillar MAIT cells were highly resistant to this toxin, and this resistance correlated with low CCR2, CXCR1 and CXCR2 expression. Notably, similarly low HlgAB receptor expression was evident in MAIT cells in barrier tissues such as lung and intestines. This is consistent with the recent findings that MAIT cells in the appendix express lower levels of CCR2 than their circulating counterparts, and this expression was further diminished in the presence of innate cytokines or the MAIT cell antigen 5-OP-RU (56). CCR2 is influenced by the local milieu and has a role in recruiting MAIT cells to inflamed tissues (56). Thus, one might speculate that CCR2 downregulation on MAIT cells at these tissue sites occurs through multiple mechanisms, including ligand-induced internalization and transcriptional suppression induced by inflammatory cytokines. Collectively, these findings indicate that MAIT cells display tissue- and context-dependent sensitivity to *S. aureus* leukocidins. These tissue-dependent patterns suggest that effective anti-staphylococcal therapies should be tailored to the anatomical site of infection and the local receptor landscape.

In summary, these findings indicate that MAIT cells respond vigorously to *S. aureus* with a diverse effector response, where the combination of functions deployed depend on the levels and kinetics of MR1-mediated antigen presentation. The broadly expressed *S. aureus* toxin HlgAB targets peripheral blood MAIT cells via their high CCR2 expression. However, MAIT cells in barrier tissues, such as the tonsil and intestine, have low CCR2 levels and are largely resistant to this toxin. Furthermore, MAIT cell activation partly protects both the MAIT cells and monocytes in the immediate vicinity. These findings provide a solid foundation for future research investigating how MAIT cells, and, more importantly, different MAIT cell subsets could be therapeutically combined with antibiotics by identifying antimicrobial pathways modulated through cytokines, cytolytic effector molecules, or chemokine receptor targeting to enhance immune responses against antimicrobial-resistant *S. aureus* infections. Importantly, such investigations should encompass tissue-specific analyses beyond blood-based studies.

## Materials and Methods

### Tissues and primary cells

Human tissue samples were obtained through the Immunology Human Organ Donor Programme (IHOPE) from deceased organ donors who had provided consent for research. Inclusion criteria followed established national guidelines for organ donation and transplantation. Tissues not used for transplantation, and biopsies from liver, were collected for research purposes. Serological screening of donors was performed according to clinical routine and included tests for cytomegalovirus, hepatitis B surface and core antigens, HIV-1, and syphilis. The following organ donor (n=9) characteristics were recorded: age (mean=58, median=62, and range=32-74 and sex (male=3, female=6). Donor metadata is available in Supplementary Table S1. The study was approved by the Swedish Ethical Review Authority.

Tonsil samples were collected from individuals undergoing tonsillectomy for recurrent infections or obstructive sleep apnea as part of the TONCIM Tissue Collection Programme at ÖNH Odenplan, Stockholm, Sweden, under approval by the Swedish Ethical Review Authority. For this analysis, 32 donors were included in total: 25 with tonsil samples only and 7 with both tonsil and PBMC samples. Donor characteristics included age (mean=27.9, median=26, range=16-55) and sex (male=18, female=14). Donor metadata is available in Supplementary Table S2. For the remaining tonsil staining fresh tonsil samples were used. Cells were plated at 2×10⁶ per well in 96-well U-bottom plates (Corning) and used for staining. For this analysis, 14 donors were included in total. Donor characteristics included age (mean=26.6, median=21, range=12-55) and sex (male=7, female=7). Donor metadata is available in Supplementary Table S3.

Peripheral blood samples were collected from healthy adult donors at Karolinska University Hospital under approval from the Swedish Ethical Review Authority. Additional detailed materials and methods descriptions relating to tissue processing and cell preparation is available in the Supplementary material.

### Antigens, microbes and cell lines

The potent MAIT cell agonist 5-(2-oxopropylideneamino)-6-(D-ribitylamino)uracil (5-OP-RU) was synthesized as a solution in DMSO-d_6_ as previously described (57, 58). *S. aureus* USA300 SF8300 was used in this study as a low-passage, PVL-positive community-associated strain representative of isolates from the United States (59). Similar to most *S. aureus* isolates, the γ-haemolysin genes are present in the core genome and SF8300 produces functional HlgAB toxin that targets chemokine receptors (34). Bacteria were cultured overnight at 37°C in casein hydrolysate and yeast extract (CCY), and bacterial counts determined with BactoBox® (SBT Instruments, Denmark) according to the manufacturer’s protocol. The microbes were then stored at -80°C in 50% glycerol/50% PBS. The THP-1 cell line (ATCC, Mananass, VA, USA) was cultured in RPMI-1640 complete medium supplemented with 25 mM HEPES, 2 mM L-glutamine (GE Healthcare), 10% fetal bovine serum (Sigma-Aldrich), 50 μg/ml gentamicin (ThermoFisher Scientific), and 100 μg/ml Normocin (Invivogen), and was routinely tested negative for mycoplasma.

### MAIT cell activation assays

Two types of activation assays were performed. In the first, MAIT cells were isolated from PBMCs and co-cultured with THP-1 cells as APCs to study MAIT cell responses to *S. aureus* (37). THP-1 cells were seeded in R10 complete medium for 2 h prior pulsing with bacteria. *S. aureus* was washed once in PBS before mild fixation in 1% formaldehyde for 3 min, followed by extensive PBS washes. The MAIT cell activating antigens present in the bacteria are not destroyed by mild fixation, and activation induced by mildly fixed bacteria is comparable to live bacteria (60). The bacteria were resuspended in complete medium and used to pulse THP-1 monocytic cells at the specified microbial dose. Subsequently, MAIT cells were added at a 2:1 ratio (MAIT:THP-1) at the indicated time points. Monensin and brefeldin A (both from BD Biosciences) were added the last 6 h of culture before staining. In selected experiments designed to evaluate whether MAIT cell responses to *S. aureus* required both MR1-mediated antigen presentation and cytokine co-stimulation, cells were treated with 20 μg/ml MR1-blocking monoclonal antibody (26.5, BioLegend) or an isotype control (IgG2a). For IL-12 and IL-18 blocking experiments, isotype control (IgG1), anti-IL-12p70 mAb (C8.6, Miltenyi Biotec) and/or anti-IL-18 mAb (125-2H, MBL Life science) were added at 5 μg/ml. All antibodies were added 3 h after the addition of the microbes. For the second assay, PBMCs were plated at 1×10⁶ per well in 96-well U-bottom plates and were stimulated with either 5-OP-RU or innate cytokines. Specifically, 5-OP-RU was added at 16 nM, or recombinant IL-12 (PeproTech) and IL-18 (MBL) were added at final concentrations of 10 ng/ml and 100 ng/ml, respectively.

### Toxin assay

PBMC or tonsillar cells were incubated with recombinant HlgAB (IBT Bioservices) at the indicated concentration between 30 min and 4 h, as indicated in the text of each figure legend. To investigate possible CCR2 antagonist-mediated protection against HlgAB toxicity, cells were pre-treated with Cenicriviroc (MedChem Express) or BMS CCR2 22 (MedChem Express) for 1 h before HlgAB addition and maintained in the medium.

### Flow cytometry

Cells were plated in 96-well plates and incubated with Fc-block (Merck) for 10 min at room temperature. Monoclonal antibodies for cell surface and intracellular staining are listed in Supplementary Table S4. For identification of MAIT cells, tetramer staining with 5-OP-RU-loaded h-MR1-PE or h-MR1-APC (NIH Tetramer Core Facility) was performed 30 min at 4°C before staining with monoclonal antibodies for other markers. If chemokine receptor staining was required, it was performed together with surface staining by incubating cells at 37°C for 30 min. In the absence of chemokine receptor staining, all other surface markers were stained at 4°C for 20 min. After staining, cells were washed with PBS containing 2% FCS and 2 mM EDTA. Intracellular staining was performed using the BD Fixation/Permeabilization Kit (BD Biosciences). Flow cytometry data was acquired on BD Symphony A5 and A3 instruments (BD Biosciences) and analyzed using FlowJo software v. 10.10 (TreeStar). For some experiments assessing toxin effects in PBMCs, BD Trucount™ Tubes were used to obtain absolute cell counts. Polyfunctionality was analyzed using SPICE v6.1 (61) and COMPASS (62), enabling visualization of individual functional subsets and Bayesian quantification of the combinatorial response.

### Statistics

Data visualization was performed in R (version 4.4.3) and GraphPad Prism v.6.0c (GraphPad Software). Statistical analyses were performed using R. Statistical comparisons between paired samples were performed using the Wilcoxon signed-rank test. When multiple pairwise comparisons were performed, p-values were adjusted using the Benjamini-Hochberg method. Comparisons of two unpaired groups were made with the Wilcoxon rank-sum test/Mann-Whitney, whereas more than two unpaired groups were analyzed using the Kruskal-Wallis and Dunn post hoc tests. Where SPICE was used, permutation test was performed with the SPICE software to compare the different groups. P-values < 0.05 were considered significant.

## Acknowledgements

The *S. aureus* USA300 SF8300 was provided by András N. Spaan and previously described (34). The MR1 tetramer technology was developed jointly by Dr. James McCluskey, Dr. Jamie Rossjohn, and Dr. David Fairlie, and the material was produced by the NIH Tetramer Core Facility as permitted to be distributed by the University of Melbourne.

## Funding

This research was supported by grants to J.K.S. from the Swedish Research Council (2023–02857), the Swedish Cancer Society (23-2912Pj), the Swedish Heart-Lung Foundation (20230551), and the Center for Innovative Medicine (FoUI-989096). E.J.M.R. was supported by a fellowship from the Wenner-Gren Foundation (UPD2022-0155). The organ donor consortium IHOPE project is supported by a grant to M.B., J.Mj. and C.J. from the Knut and Alice Wallenberg Foundation (KAW 2022.0021). D.P.F. was supported by grants from the Australian National Health and Medical Research Council (2009551), the US National Institutes of Health (R01 AI14807-01A1), and the Australian Research Council (CE200100012). N.M. was supported by the Swedish Research Council (2021–03069) and the Swedish Cancer Society (22-2319Pj). The funders had no role in study design, data collection and analysis, decision to publish, or preparation of the manuscript.

## Ethics declaration

The study was approved by the Swedish Ethical Review Authority (IDs: 2020-02-604; 2024-04466-2; 2021-03694). and performed in accordance with the Declaration of Helsinki.

## Conflict of interest statement

The authors have declared that no conflict of interest exists.

## Authorship contributions

E.J.M.R. and J.K.S. designed the study.

E.J.M.R and J.T. performed experiments.

C.B., E.M. and E.L. developed methodology.

E.M., C.Ca., T.K., V.N., T.S., C.Co., E.W., T.R.M., A.Mi., A.Ma., C.T., S.F., J.K., M.F., J.B., C.S., N.W., E.B., M.E., J.D., contributed to tissue collection, processing and data acquisition.

J.Y.W.M. and D.P.F. provided critical resources.

J.K.S., N.M., A.N.T., J.M., C.J. and M.B. provided supervision and funding for the study.

E.J.M.R. and J.K.S. interpreted the data and wrote the paper.

All authors read and revised the manuscript.

## Appendix

Supplementary material and methods, supplementary tables 1-4 and supplementary figures 1-3 are available in the supplementary items file.

